# REM sleep: unique associations with behavior, corticosterone regulation and apoptotic pathways in chronic stress in mice

**DOI:** 10.1101/460600

**Authors:** Mathieu Nollet, Harriet Hicks, Andrew P. McCarthy, Huihai Wu, Carla S. Möller-Levet, Emma E. Laing, Karim Malki, Nathan Lawless, Keith A. Wafford, Derk-Jan Dijk, Raphaelle Winsky-Sommerer

## Abstract

One of sleep’s putative functions is mediation of adaptation to waking experiences. Chronic stress is a common waking experience, however, which specific aspect of sleep is most responsive, and how sleep changes relate to behavioral disturbances and molecular correlates remain unknown. We quantified sleep, physical, endocrine and behavioral variables and the brain and blood transcriptome in mice exposed to nine weeks of unpredictable chronic mild stress (UCMS). Comparing 46 phenotypical variables revealed that rapid-eye-movement sleep (REMS), corticosterone regulation and coat state were most responsive to UCMS. REMS theta oscillations were enhanced whereas delta oscillations in non-REMS were unaffected. Transcripts affected by UCMS in the prefrontal cortex, hippocampus, hypothalamus and blood were associated with inflammatory and immune responses. A machine learning approach controlling for unspecific UCMS effects identified transcriptomic predictors for specific phenotypes and their overlap. Transcriptomic predictor sets for the inter-individual variation in REMS continuity and theta activity shared many pathways with corticosterone regulation and in particular pathways implicated in apoptosis, including mitochondrial pathways. Predictor sets for REMS and anhedonia, one of the behavioral changes following UCMS, shared pathways involved in oxidative stress, cell proliferation and apoptosis. RNA predictor sets for non-NREMS parameters showed no overlap with other phenotypes. These novel data identify REMS as a core and early element of the response to chronic stress, and identify apoptotic pathways as a putative mechanism by which REMS mediates adaptation to stressful waking experiences.

**Significance Statement:** Sleep is responsive to experiences during wakefulness and is altered in stress-related disorders. Whether sleep changes primarily concern rapid-eye-movement sleep (REMS) or non-REM sleep, and how they correlate with stress hormones, behavioral and transcriptomic responses remained unknown. We demonstrate using unpredictable chronic (9-weeks) mild stress that REMS is the most responsive of all the measured sleep characteristics, and correlates with deficiency in corticosterone regulation. An unbiased machine learning, controlling for unspecific effects of stress, revealed that REMS correlated with RNA predictor sets enriched in apoptosis including mitochondrial pathways. Several pathways were shared with predictors of corticosterone and behavioral responses. This unbiased approach point to apoptosis as a molecular mechanism by which REMS mediates adaptation to an ecologically relevant waking experience.

## Introduction

Sleep is assumed to contribute to recovery from the wear and tear of wakefulness and to mediate adaptation to the waking experience, be it through memory consolidation or processing of emotional experiences such as those associated with stressful events (1). Chronic stress is the most significant predictor of mood disorders (2) and major depressive disorder is anticipated to be the leading cause of disease burden by 2030 (3), while the true global burden of stress-related mental diseases might be largely underestimated (4).

In animals, chronic stress leads to profound physiological changes, such as hypothalamic-pituitary-adrenal (HPA) axis regulation of corticosterone, neurogenesis, synaptic plasticity and gene expression (5-7). Chronic stress also leads to a plethora of behavioral disturbances including depressive-like behavior, decreased responsiveness to rewards akin to anhedonia, a core symptom of depression, and sleep alterations (1, 8). The effects of chronic stress on sleep in rodents have been studied by applying physical, social and/or environmental stressors. Several of these studies documented alterations in rapid eye movement sleep (REMS) and sleep continuity (9-12), while others reported changes in non-REM sleep (NREMS) and the electroencephalogram (EEG) slow wave (delta) activity (13). In humans, chronic stress, alterations in the HPA axis regulating cortisol and sleep disturbances have been associated with mood disorders (14, 15). However, the nature of sleep disturbances in major depression continues to be discussed, with some studies highlighting changes in NREMS including alterations in slow wave (or delta) oscillations (16, 17), and others REMS and sleep continuity (15, 18). Unresolved questions are how the various physiological and behavioral consequences of chronic stress interrelate and whether specific changes in sleep are early and core symptoms contributing to adaptation to chronic stress.

Altered patterns of gene expression in the brain are triggered by stress and transcriptome responses have been shown to be highly tissue/brain region specific (6). Most studies have focused on the hippocampus and prefrontal cortex, identifying differential gene expression in processes related to inflammation, immune response and neurogenesis (19-21). While transcriptomic changes underlying neuroplastic adaptation to chronic stress have been extensively studied, very few animal studies investigated the transcriptome response to stress in blood (22, 23). This is of particular interest in the context of translational studies since blood transcriptomic signatures of depression and treatment response have been identified in humans (24-26). Finally, the extent to which sleep and other behavioral and endocrine alterations in response to stress are related to changes in the transcriptome has not yet been comprehensively quantified.

Here, exposure to chronic stress was achieved using the well-validated unpredictable chronic mild stress (UCMS) paradigm in mice (8). UCMS elicits a broad range of physiological and ethological changes which are consistent with symptoms of major depressive disorder, and predicts the efficacy of antidepressant treatments (8, 27). This ethological ‘model’ has been recognized for its high translational potential in the context of stress-related disorders (28, 29). The aims of the current study were 1) to comprehensively characterize changes in REMS and NREMS, corticosterone and behavioral variables, as well as the transcriptomic signature in three stress- and sleep-related brain regions (hippocampus, prefrontal cortex, hypothalamus) and blood, in response to chronic stress and 2) use machine learning and other robust statistical approaches to investigate the interrelationship of these responses.

## Results

### Stress-induced physical, neuroendocrine and behavioral disturbances

We assessed the impact of the repeated exposure to an unpredictable stressful waking experience on a number of physiological and behavioral variables during the 9-week protocol (Fig. 1A). Chronic mild stress significantly altered body weight and worsened coat state, an index of reduced grooming behavior (Fig. 1B-C). The corticosterone regulation was compromised in the UCMS group, consistent with blunted HPA axis negative feedback (Fig. 1D). Self-care behavior was reduced, as reflected by increased grooming latency and decreased grooming duration (Fig. 1E-F). Quality of nest building, indicative of motivation, was also reduced in the UCMS group (Fig. 1G). Moreover, UCMS suppressed the progressive increase of consumption of a palatable stimulus, indicative of anhedonia (Fig. 1J-K). Immobility during the forced swim test was increased (Fig. 1L), as was anxiety-like behavior (Fig. 1M). Social disturbances were observed with increased aggressive behavior, and decreased social preference for the novel congener (Fig. 1N-O). Exposure to UCMS reduced the weekly averaged locomotor activity during the dark (active) phase of the light-dark cycle, while activity remained unaffected during the light phase (Fig. 1H-I). A secondary analysis of the 24-h time course showed that the UCMS group was less active in the dark phase during the last three weeks of the 9-week UCMS paradigm (*SI Appendix*, Fig. S1; for detailed statistics, see *SI Appendix*, Datasets S1).

**Fig. 1.**
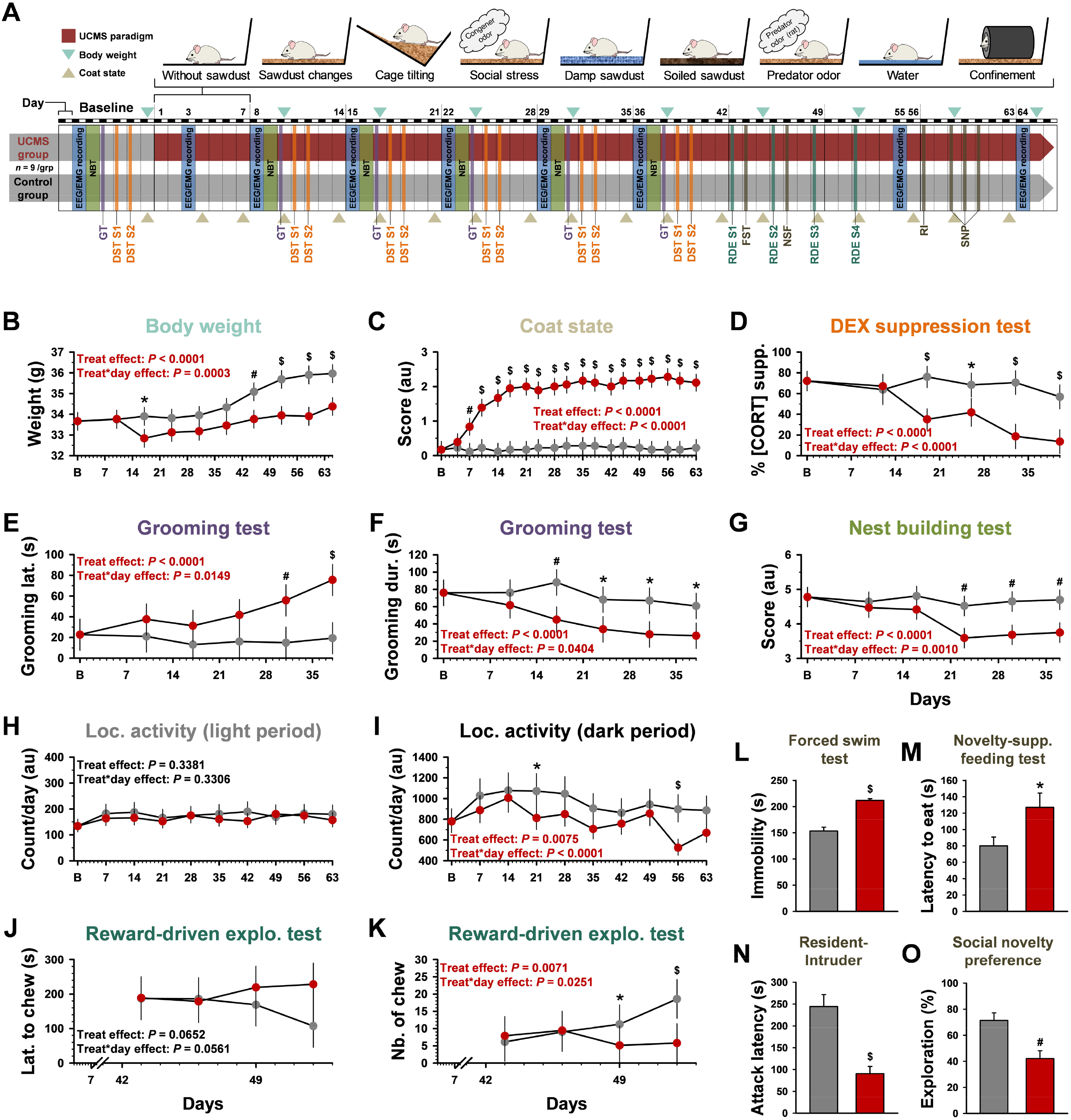
Unpredictable Chronic Mild Stress (UCMS) protocol and induced physical, corticosterone regulation and behavioral alterations. (A) Overview of the protocol. Mice were randomly assigned to the control (grey) or the UCMS (red) group. (B) Body weight, (C) coat state, a surrogate of self-care behavior, (D) hypothalamic-pituitary-adrenal axis negative feedback assessed at the daily peak of corticosterone [dexamethasone (DEX) suppression test; DST; *n* = 5-7 per group], (E-F) self-centered behavior (grooming test; GT: Grooming Latency and Grooming Duration), (G) motivation (nest building test; NBT), and (H-I) locomotor activity, were measured at baseline and during the nine-week UCMS. From day #43, several behavioral domains were evaluated, including (J-K) anhedonia-like (reward-driven exploratory test assessing the motivation to collect a palatable stimulus; Latency to chew and Number of Chews; (L) despair (depressive-like) behavior assessed by increased immobility in the forced swim test (FST; *n* = 8 per group), (M) anxiety-like behavior evaluated by increased latency to eat the food pellet in the novelty-suppressed feeding test (NSF), (N) aggressiveness identified by shorter attack latency in the resident-intruder (RI) test and (O) social disturbances reflected by reduced time spent with the unfamiliar conspecific in the UCMS group (social novelty preference test; SNP). Data are shown for *n* = 9 per group (unless specified otherwise), as Least-Squares Mean ± 95% Confidence Intervals, except for (L-O) Mean ± SEM; ^*^*P* < 0.05, ^#^*P* < 0.01, ^$^*P* < 0.001 (*post-hoc* comparisons for significant ‘treatment’ x ‘day’ interaction in general linear mixed model, or significant *t*-test for non-repeated measures). S: session.

### Impact of a stressful experience on sleep structure, sleep continuity and the spectral composition of the electroencephalogram

We also assessed the effect of 9-week UCMS on sleep and the EEG. Twenty-four-hour total sleep time (TST) and NREMS duration were not significantly altered during UCMS (TST: ‘treatment’ effect *P* = 0.4727; interaction ‘treatment’ x ‘day’: *P* = 0.0993; NREMS: ‘treatment’ effect *P* = 0.5088; interaction ‘treatment’ x ‘day’: *P* = 0.1898; *SI Appendix*, Datasets S1). By contrast, a significant increase in 24-h REMS duration was observed (‘treatment’ effect *P* < 0.0001; interaction ‘treatment’ x ‘day’: *P* < 0.0001; *SI Appendix*, Datasets S1). Expressed as a percentage of TST, REMS was increased while NREMS decreased (Fig. 2A-2D), and these changes were observed both during the light and dark phases (*SI Appendix*, Fig. S2A-D). Analysis of the 24-h time course of REMS confirmed an overall treatment effect throughout the UCMS protocol (*SI Appendix*, Fig S3F-J), while the 24-h time course of TST and NREMS revealed a ‘treatment’ x ‘time of day’ interaction during the second half of the protocol with an increased sleep duration primarily during the dark phase (*SI Appendix*, Fig. S3A-E and S3P-T). Chronic mild stress also induced an overall increase in REMS continuity, with increased number and duration of REMS episodes (Fig. 2B-C) during both the light and dark phases (*SI Appendix*, Fig. S2G-H and S2K-L). By contrast, NREMS became more fragmented with an increased number of episodes of shorter duration (Fig. 2E-F and *SI Appendix*, Fig. S2E-F and S2I-J; for detailed statistics, see *SI Appendix*, Datasets S1).

**Fig.2.**
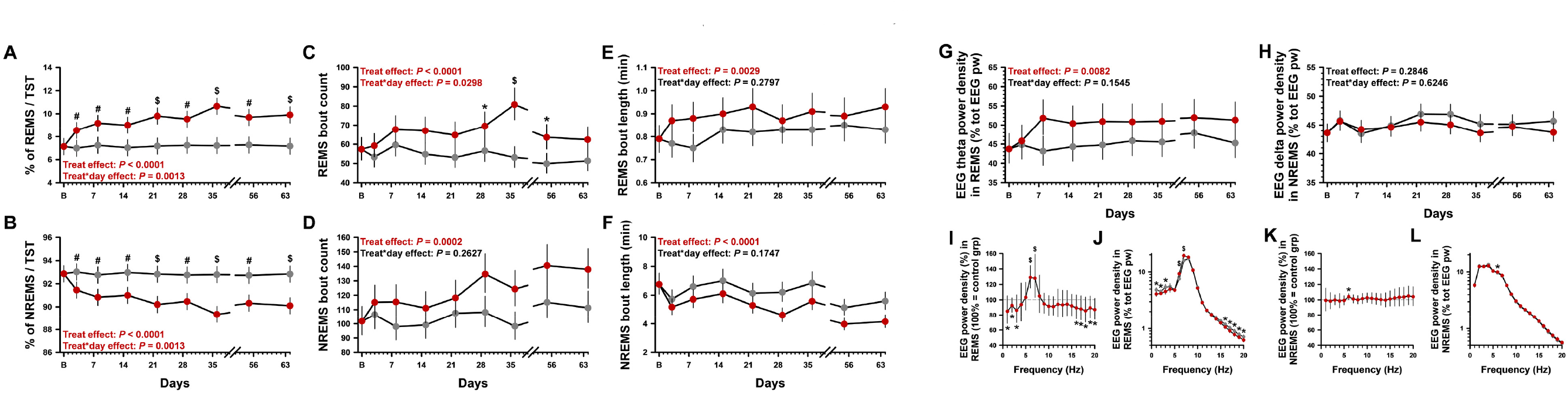
Time-course of UCMS-induced alterations on sleep and the electroencephalogram (EEG). (A-B) Rapid-eye movement sleep (REMS) and non-rapid eye movement sleep (NREMS) as percentage of total sleep time per 24-h (TST). (C-D) Number and (E-F) duration of sleep episodes; (G) EEG theta power density (6-9 Hz) in REMS and (H) EEG delta activity (1-4.5 Hz) in NREMS during the 12-h light phase. (I-L) Relative EEG power spectra in REMS and NREMS during the light phase. Data are Least-Squares Mean ± 95% Confidence Intervals (controls: grey; UCMS: red; *n* = 8 per group); \**P*<0.05, #*P*<0.01, $*P*<0.001 (*post-hoc* comparisons for (A-H) significant ‘treatment’ x ‘day’ interaction or (I-L) effect of ‘treatment’ in general linear mixed model). NREMS: non-rapid eye movement sleep; grp: group; pw: power; REMS: rapid eye movement sleep; tot: total.

Quantitative EEG analysis, using baseline measurements as a covariate to control for individual differences in the EEG spectra, showed that theta activity, an EEG hallmark of REMS, was increased in the light (Fig. 2G) and dark phases (*SI Appendix*, Fig. S2O). By contrast, NREMS delta activity was not affected by UCMS (Fig. 2H and *SI Appendix*, Fig. S2). Computation of relative EEG power spectra showed that changes in REMS were indeed mainly observed in the theta range, although some reduced activity in lower and higher frequencies was detected (Fig. 2I-J). Only minor changes were observed in the relative NREMS EEG power spectra (Fig. 2K-L). The 24-h time course of EEG theta power showed no significant interaction of ‘treatment’ x ‘time of day’ (*SI Appendix*, Fig. S3K-O), while EEG delta power was only reduced during the second part of the dark phase and only during the last week of the protocol (*SI Appendix*, Fig. S3U-Y; for detailed statistics, see *SI Appendix*, Datasets S1).

### Temporal associations of physical, neuroendocrine, behavioral and sleep disturbances

The changes in 24-h REMS duration, and other measures of sleep duration across 24h or during the light phase, were observed as early as day #3 of the UCMS protocol (Fig. 2A and *SI Appendix*, Fig. S4 and Fig. S2A-D). Degradation of coat state occurred from day #7, while differences in body weight, impairment of corticosterone regulation, self-centered behavior and motivation appeared in weeks #3-4 (Fig. 1B-G and *SI Appendix*, Fig. S4). Locomotor activity in the dark period was altered only towards the end of the 9-week protocol (*SI Appendix*, Fig. S1B-E).

### Effect size and stability of chronic stress effects across physical, neuroendocrine, behavioral and sleep symptoms

The size of the effects of UCMS varied considerably across dependent measures, with the largest effect sizes observed for coat state, 24-h REMS duration, corticosterone regulation and 24-h REMS expressed as percentage of TST (Fig. 3A). Overall, most REMS and NREMS variables, including the number and length of sleep episodes, displayed a large or medium effect size (Cohen’s *f*2 > 0.25, Fig. 3A). Across behaviors, effect sizes of UCMS were large for despair behavior, aggression, self-centered behavior, social disturbances, anxiety-like behavior and motivation. The impact of UCMS on 24-h TST, 24-h NREMS duration and EEG delta power was small (Fig. 3A).

**Fig.3.**
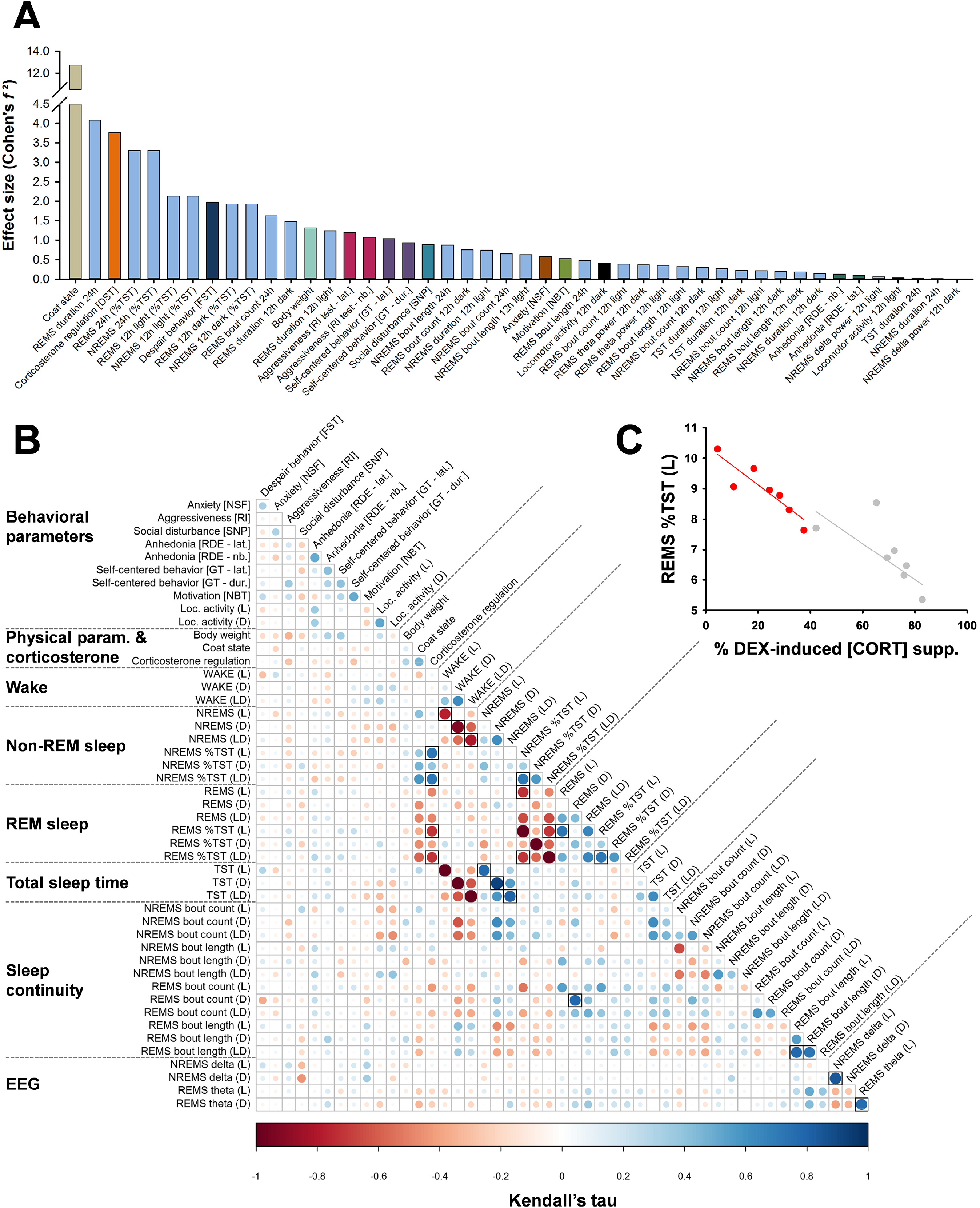
Effect size and phenotypic interactions of UCMS-induced physical, behavioral, neuroendocrine, neurochemical and sleep-related symptoms. (A) Effect sizes (Cohen’s *f^2^*) of repeated and non-repeated measures (large effect size: > 0.40; medium: 0.25 to 0.40; small: 0.10 to 0.25). Effect sizes computed for non-repeated measures (Cohen’s *d*) were converted to Cohen’s *f^2^* Figures and Table using the following formula: Cohen’s *d* = 2 x Cohen’s *f2*. (B) Kendall’s partial correlation between pairs of phenotypes (i.e., after removing the effect of the experimental groups). The phenotypes (the averaged last three measurements were used for repeated measures) were ordered according to their phenotypic categories. Correlations were considered significant at a false discovery rate (FDR) < 0.05 (*P*adj; symbolized by black square), computed with the Benjamini-Hochberg procedure for multiple testing correction. (C) Example of a correlation from (B) illustrated for REMS percentage during the light (L) phase and impairment of the corticosterone regulation (*P*_nom_=0.00034, *P*_adj_= 0.0197; *n* = 7 animals per group; grey: control mice, red: UCMS-subjected animals). DEX supp: dexamethasone suppression.

In addition, to assess to which extent UCMS-induced changes were stable within individuals, intra-class correlation (ICC) coefficients were computed for all dependent variables. ICCs ranged between 0.67 and 0.997 for body weight, locomotor activity, REMS EEG theta power and NREMS delta power, suggesting that the response to UCMS is highly stable within individuals (benchmarks defined by (30)). Coat state, and REMS and NREMS expressed as a percentage of TST for 24-h showed moderate trait stability (ICC = 0.5240 and 0.4671 respectively). Corticosterone regulation displayed a ‘slight’ stability (ICC = 0.0066) (see *SI Appendix*, Datasets S1).

### Bivariate associations across physical, behavioral, sleep and neuroendocrine disturbances

To assess the strength of associations between measured variables, we computed Kendall’s partial correlations between pairs of symptoms induced by chronic mild stress. We controlled for the effect of ‘group’ (i.e., control *versus* UCMS) to identify bivariate associations at the level of the individual independent of ‘unspecific’ group effects. The increased percentage of REMS per TST (during the light phase and for 24-h) correlated negatively with dexamethasone suppression, i.e., more REMS was associated with the impairment of corticosterone regulation (Tau = 0.72, nominal *P* value (*P*_nom_) = 0.00034, False Discovery Rate-adjusted *P* value (*P*_adj_) = 0.0252 and Tau = 0.71, *P*_nom_ = 0.00037, *P*_adj_ = 0.0211, respectively; Fig. 3B-C and *SI Appendix*, Datasets S2). These associations were of a large effect size defined by Tau > 0.25 (31).

### Effects of 9-week chronic stress on the transcriptome of prefrontal cortex, hippocampus, hypothalamus and blood

To gain further insight into the molecular mechanisms underlying the phenotypes induced by UCMS, we performed RNA-Sequencing on brain region and whole blood samples collected at the end of the UCMS paradigm.

### Differential gene expression and functional enrichment

We first assessed differential expression between the UCMS and control groups. The number of differentially expressed genes (DEGs) was relatively small (*SI Appendix*, Fig. S5). The ratio between up-regulated and down-regulated genes was positive for all tissues (*SI Appendix*, Fig. S5). A literature search revealed that numerous DEGs had been previously reported to be associated with sleep and/or circadian rhythms, stress, neuropsychiatric symptoms, mood disorders or neurodegenerative diseases such as Alzheimer’s and Parkinson’s diseases (Fig. 4A and *SI Appendix*, Datasets S3).

**Fig.4.**
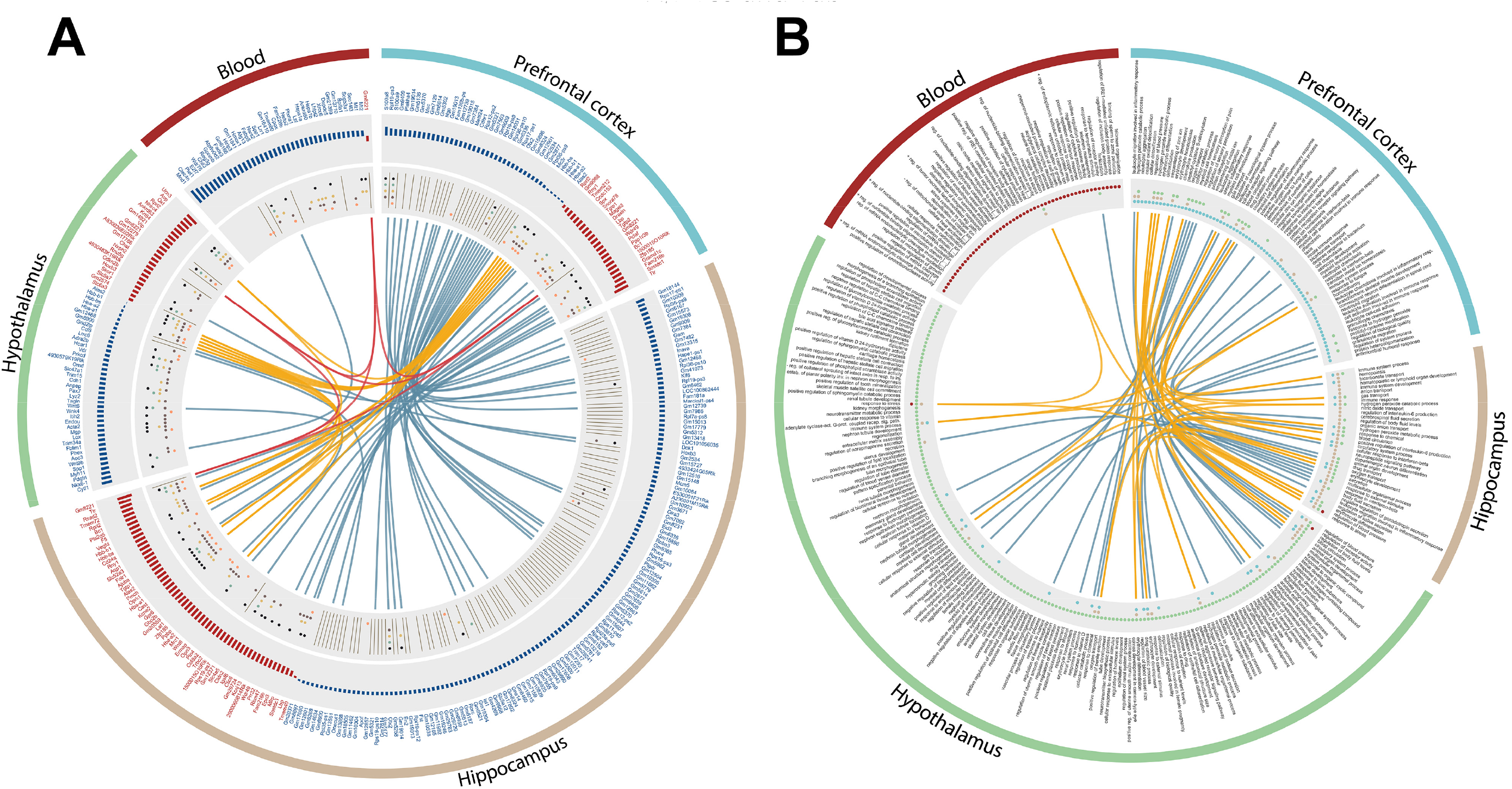
Characterization and functional enrichment of genes differentially expressed following chronic mild stress in different tissues. (A) Differentially expressed genes. The various tracks/circles indicate the following. Outer track: Tissue. Second track: Gene names: blue upregulated genes (*P_pfp_*<0.05). Red: downregulated genes (*P_pfp_*<0.05). Third track: length of blue and red bars indicate log_2_ fold-change of the DEGs. Innermost grey track: The dots refer to conditions in which the DEGs have been reported as being associated with/implicated in. Conditions: stress response (brown dots; sleep (orange dots); behavioral and neurobiological changes corresponding to human ‘neuropsychiatric’ symptoms (ochre); mood disorders (green) and neurodegenerative diseases (black) [for references, see *SI Appendix*, Datasets S3]. The grey lines in the innermost grey track indicate processed pseudogenes. The arcs in the center circle depict the overlap between tissues. Blue arc: two tissues. Orange arc: three tissues; Red arc: four tissues. (B) Significantly enriched Gene Ontology (GO) biological processes for differentially expressed genes (DEGs) in the prefrontal cortex, hippocampus, hypothalamus and whole blood, after nine-week unpredictable chronic mild stress. Enrichment analyses were performed using MetaCore™ ‘GO Processes’ (GeneGo, Thomson Reuters; https://portal.genego.com/) and significance was set at FDR adjusted *P*-value (*P*_adj_) < 0.05. The various tracks/circles indicate the following: Outer track: Tissue. Second track: processes names. Third track: tissues in which biological processes were found to be significantly enriched (blue: prefrontal cortex; brown: hippocampus; green: hypothalamus; red: blood). Overlaps between the different tissues is illustrated by arcs in the inner circular track (blue: overlap between two tissues; yellow: overlap between three tissues); *n* = 8 per group for brain regions; *n* = 7 controls *versus n* = 9 UCMS group for blood. This information is available in tabular format (see *SI Appendix*, Datasets S4).

In the prefrontal cortex, at the individual transcript level, *S100a8* and *S100a9,* playing a key role in the regulation of the inflammatory and immune responses, were among the most up-regulated protein-coding DEGs (Fig. 4A and *SI Appendix*, Datasets S3). *Mc3r*, encoding a receptor for melanocyte-stimulating hormone and adrenocorticotropic hormone and involved in the regulation of the circadian clock, was also up-regulated. The most down-regulated cortical genes included *Ttr*, encoding transthyretin, thyroid hormone-binding protein, a carrier protein also involved in amyloid deposition, and genes involved in the innate immune system or inflammatory response (*Lbp; Tgtp2*) (Fig. 4A and *SI Appendix*, Datasets S3). To further characterize the effects of the 9-week UCMS, we next performed functional enrichment using Gene Ontology (GO) processes and canonical pathway maps. DEGs were associated with seventy-four significantly enriched biological processes in the prefrontal cortex, including several processes involved in inflammatory and immune response, metabolic processes and signaling pathways among others (Fig. 4B and *SI Appendix*, Datasets S4). Enriched pathways evoked by chronic stress were involved in transcription with the Role of AP-1 in regulation of cellular metabolism and development (*SI Appendix*, Fig. S6 and Datasets S4). In the hippocampus, the most up-regulated protein-coding gene was *Gem,* possibly involved in receptor-mediated signal transduction and previously associated with sleep deprivation (Fig. 4A and *SI Appendix*, Datasets S3). Other up-regulated genes included *Inava*, encoding the innate immunity activator protein, *Glra3* encoding the glycine receptor alpha3 subunit, *Ptgdr* encoding the prostaglandin D2 receptor, and *Rorc* encoding the nuclear receptor ROR-gamma a key regulator of immunity and peripheral circadian rhythms. One of the most down-regulated genes was *Ttr*, as observed in the prefrontal cortex. Other hippocampal down-regulated DEGs were involved in the immune system, inflammation and the vascular system (e.g*., Lbp*; *Rsad2; Pla2g5*; *F5; Vegdf; Cd24a; Tgtp1; Cast*; *Lst1*), circadian rhythms (*Opn3*), and neural transmission (e.g., *Pmch*, *Oprk1*, *Kcne2*, *Gpr6*). Response to external stimulus, cerebrospinal fluid secretion and cellular response to interferon-beta were among the enriched GO biological processes in the hippocampus (Fig. 4B and *SI Appendix*, Datasets S4). Enriched pathway maps induced by UCMS were linked to transcription, development and immune response. However, none were significant after FDR adjustment (*SI Appendix*, Fig. S6 and Datasets S4).

In the hypothalamus, some of the up-regulated protein-coding genes induced by UCMS are involved in the Wnt/β-Catenin signaling interactive pathway, cell-matrix interaction and cell adhesion (e.g., *Wnt9b, Wnt6, Spp1, Aoc3, Fblim1, Mgp, Acta2, Itih2, Cdh1*) which have also been previously implicated in stress response (Fig. 4A and *SI Appendix*, Datasets S3). Among the most down-regulated genes in the hypothalamus were DEGs encoding dopamine and choline transporters (*Slc6a3, Slc5a7*), as well as *Chat*, encoding Choline O-acetyltransferase that catalyzes the biosynthesis of acetylcholine. Moreover, several DEGs encoding neuropeptide precursors involved in adaptation to stress and social behavior (i.e., *Ucn3*, *Avp*, *Oxt*, *Vip*) were down-regulated. Functional enrichment identified a large number of significant GO biological processes (n = 168), such as neurotransmitter biosynthetic processes, G-protein coupled receptor signaling pathway, response to external stimulus, grooming and aggressive behaviors (Fig. 4B and *SI Appendix*, Datasets S4). One enriched pathway, involved in protein folding and maturation and the posttranslational processing of neuroendocrine peptides, was observed (*SI Appendix*, Fig. S6 and Datasets S4).

All DEGs in blood were up-regulated except for the predicted gene *Gm8221*. Among the most up-regulated protein-coding genes, some were involved in DNA damage response (i.e., *Mnd1*; *E2f7*), the immune response and inflammation (e.g., *Clec4n*; *Chil3*; *Reg3g, Bpifa1*). Some others DEGs have been previously associated with sleep deprivation or sleep fragmentation (*Fads3*, *Gm6166, Spp1*, *Hspa1a, Hspa1b, Scgb3a1*) (Fig. 4A and *SI Appendix*, Datasets S3). Cellular processes in whole blood included regulation of cytokine production, several processes involved in RNA cleavage, unfolded protein response, nitric oxide-mediated signal transduction, and tumor necrosis factor-mediated signaling pathways (Fig. 4B and *SI Appendix*, Datasets S4).

The comparison of transcriptomic responses in the four tissues showed a robust overlap of DEGs between the prefrontal cortex and the hippocampus, while the commonalities between other tissues were weaker (Fig. 4A and *SI Appendix*, Datasets S3). The three brain regions had only six common DEGs, encoding hemoglobin subunits (*Hba-a1*, *Hba-a2*, *Hbb-b1*, *Hbb-bs*), an erythroid-specific mitochondrially located enzyme (*Alas2*), as well as the non-coding RNA *Rprl2*. Only one DEG, the predicted gene *Gm8221* (apolipoprotein L 7c pseudogene), was common to all four tissues and was among the most down-regulated DEGs in all tissues (Fig. 4A and *SI Appendix*, Datasets S3).

Several enriched biological processes were shared across two or three brain regions, such as immune system process, circulatory system process, erythrocyte development and differentiation, and oxygen transport in the prefrontal cortex, hippocampus and hypothalamus (Fig. 4B). By contrast, only two enriched GO processes were common to blood and brain regions, with response to stress in blood, hypothalamus and hippocampus and regulation of receptor activity between blood and the hypothalamus (Fig. 4B).

### Bivariate correlations between molecular consequences of chronic stress and physical, behavioral, sleep and neuroendocrine disturbances

To identify associations between differentially expressed genes and phenotypic alterations observed following 9-week UCMS, we performed bivariate analyses, computing Kendall’s partial correlations in which the effect of ‘group’ (i.e., control *versus* UCMS) was controlled for, for all physical, neuroendocrine, behavioral, and sleep variables and DEGs per tissue. When focusing on correlations of large effect size (i.e., Tau > 0.25), we observed multiple associations between DEGs and stress-induced symptoms, and in particular with sleep parameters in all four tissues (*SI Appendix*, Datasets S5). However, no correlation remained significant after FDR adjustment.

### Selection of transcriptomic predictor sets associated with complex phenotypes using a penalized regression approach

While univariate approaches provide some insights into the associations between transcripts and other physiological and behavioral variables, they nevertheless suffer from the multiplicity problem. In addition, they are not necessarily best suited to identify sets of transcripts that predict specific complex phenotypes. Thus, we next applied elastic-net learning, a multivariate approach based on a generalized linear model using penalized regression, to identify sets of features predicting specific phenotypes. We performed this analysis using all transcripts identified by RNA sequencing, i.e., not just the DEGs, focusing on sleep variables and some variables associated with stress and mood disorders. We aimed to identify transcriptomic features that were specifically associated with the sleep and behavioral variables independent of the ‘group’ effect (i.e., control *versus* UCMS). We therefore applied normalization procedures which prevented identifying features related to the unspecific effects of UCMS (see Materials and Methods). The number of features in the various identified predictor sets was overall small and varied between variables and across tissues (range: 19-333; *SI Appendix*, Datasets S6). Only very few DEGs, which are likely to primarily reflect group effects, were part of the features identified by elastic-net. To gain insights into molecular mechanisms associated with a given sleep or behavioral variable and to contrast biological correlates of the sleep and other variables, we then performed functional enrichment of predictor sets focusing on pathway maps.

### Predictor sets and functional enrichment for REMS and NREMS

While the size of the predictor sets for REMS and NREMS parameters were similar, we only observed very few significant overlaps. Common predictors were seen in the prefrontal cortex, primarily between REMS continuity and NREMS duration and continuity, as well as in the hippocampus between REMS bout count and NREMS duration and bout count (*SI Appendix*, Datasets S7). Looking individually at transcriptomic features for REMS variables, several were involved in the regulation of NF-kappa-B signaling (duration: *Otud7b*; bout-length: *Tceal7*; bout-count: *Aebp1*, *Smad6*; EEG-theta: *Card14*, *Birc3, Tceal7*). Predictors for REM duration included hippocampal transcripts playing a key role in development (*Dlx5, Tbx3, Tet3*) and/or stem cell proliferation such as *Shh* encoding sonic hedgehog, while the cold-induced mRNA binding protein coding gene *Rbm3* was a predictor of REMS duration in blood. Some sleep and circadian-related transcripts were predictors of REMS continuity and EEG theta power, including *Kcnj12, Kcnj13*, *Ciart and Ahr*. In addition, several transcripts associated with neural transmission were part of the predictor sets of REMS continuity (*Nt-3*; *Drd2*, *Mpdz*) and EEG theta power (*Htr2b; Nr4a2*). Several circadian-related transcripts were also among the predictors for NREMS (Duration: *Ppargc1a, Rorc*. Bout length: *Ciart, N1rd1*. Bout count: *Nr1d1*, *Mapk10*), as well as transcripts involved in neural transmission (Duration: *Zdhhc17*; Bout-length: *Kcnj13*, *Htr4, Ptger1*, *Pebp1*. Bout-count: *Kcnj13*, *Snap25, Lrrtm2, Pebp1*). In addition, many predictors of NREMS were associated with mitochondrial function (Duration: *Pet117, Coa7*, *Mettl20, Suclg2, Ppargc1a*, *Agk*, *Tfb2m, Immt*. Bout count: *Mrpl41, Nt5m*, *Vars2, Nr1d1, Bnip1*, *Fam210b, Tigar, Ndufaf8, Nr1d1, Slc30a10, Pars2*, *Cycs, Slc25a4*, Bout length: *Mrpl10, Mtus1, Ak2, Phb2, Sdhaf3, Pm20d1*, *Nr1d1*).

To identify whether predictors of REMS and NREMS parameters were involved in specific molecular pathways, we performed functional enrichment. Whereas REMS predictor sets were significantly enriched for many canonical pathways (n = 193), only three pathways were identified with NREMS predictor sets (Fig. 5 and *SI Appendix*, Datasets S8). Enriched pathways associated with predictors of REMS duration and REMS theta power were primarily found in the hippocampus, while pathways associated with predictors of REMS bout length and REMS bout count were enriched in the prefrontal cortex, hippocampus and hypothalamus, and in the hippocampus and blood respectively (Fig. 5). Across REMS variables and tissues, several pathways were involved in the immune response (including several Interleukin, Interferon, and Toll-like receptor signaling pathways), apoptosis and survival (e.g., Tumor Necrosis Factor [TNF] and NF-kappaB signaling pathway, role of inhibitor of apoptosis proteins, regulation of apoptosis by mitochondrial proteins, endoplasmic reticulum stress response pathway). Some enriched pathways were also associated with stem cells and development (Fig. 5). Remarkably, eleven enriched pathways were common to REMS theta power and REMS bout length, with eight of them associated with apoptosis and cell survival (some of these pathways are listed in the generic category in Fig. 5 and in Table 1). Thus, communality is non-trivial because power is a density measure which does not necessarily increase with bout duration. Of the very few significantly enriched pathways identified for NREMS predictor sets, two pathways overlapped with REMS continuity variables (Fig. 5).

**Fig.5.**
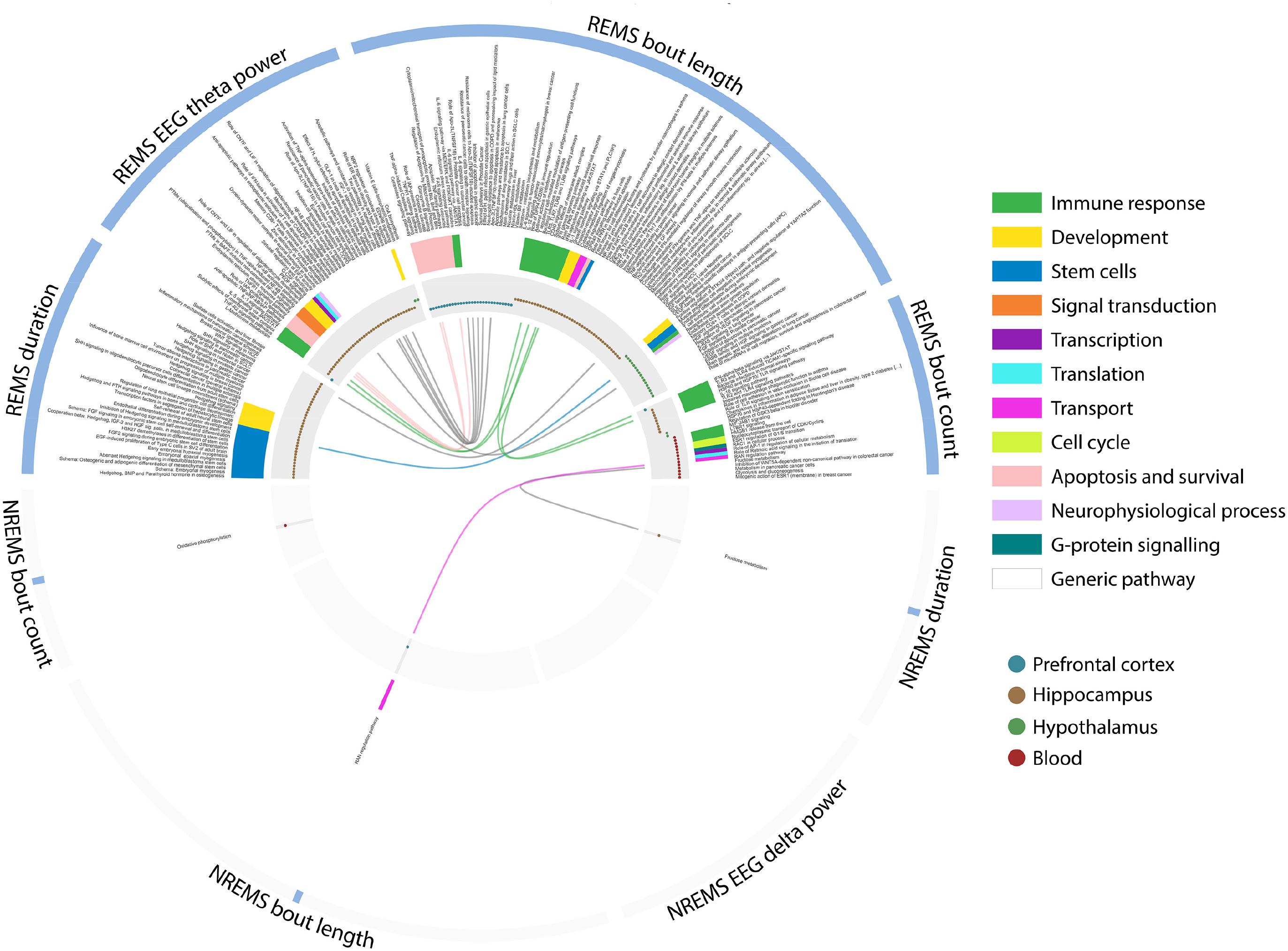
Biological pathways associated with transcriptomic predictors of sleep variables. Circular plots show enriched REMS- (top) and NREMS- (bottom) associated pathways. Tracks from outside-in, outer track: phenotypes; second track: significantly enriched pathways (identified using the ‘Pathway Maps’ tool in MetaCore™, Thomson Reuters); third track: biological categories of enriched pathways; fourth track: tissues in which pathways were found to be significantly enriched (blue: prefrontal cortex; brown: hippocampus; green: hypothalamus; red: blood). Overlaps between pathways are illustrated by arcs in the inner circular track, which color corresponds to biological categories. All depicted pathways were significant at FDR adjusted *P*-value (*P*_adj_) < 0.05; *n* = 8 per group for brain regions; *n* = 7 controls *versus n* = 9 UCMS group for blood. This information is available in tabular format (see *SI Appendix*, Datasets S8).

**Table 1.**
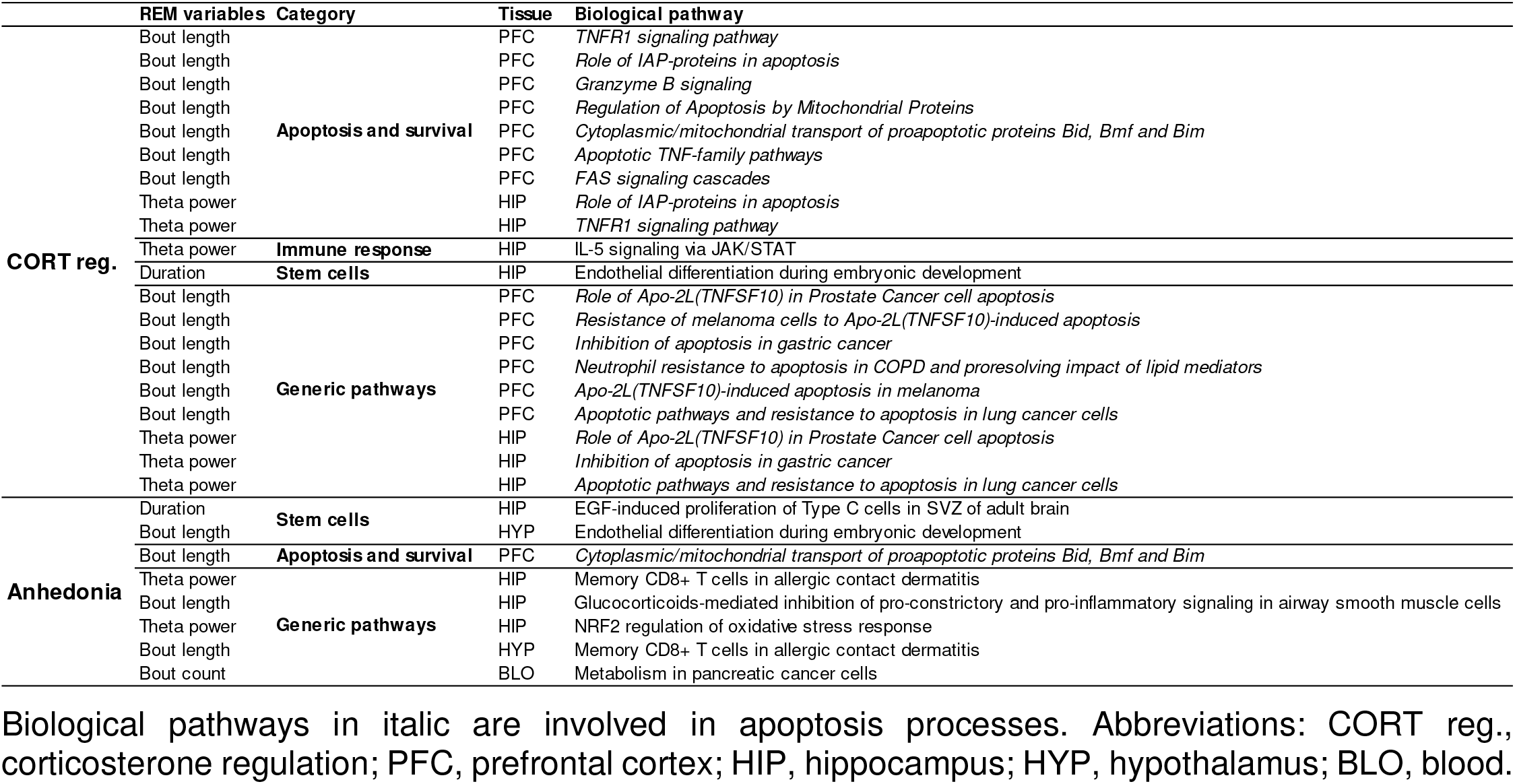
Overlap of predictor-related pathways associated with REMS parameters, corticosterone regulation and anhedonia REM variables Category Tissue Biological pathway

### Predictor sets and functional enrichment for corticosterone regulation, anxiety-, anhedonia-like and despair behaviors

Looking at individual features for the corticosterone regulation, several cortical features were associated with neural transmission and have been associated with psychiatric conditions. This includes *Cntnap4*, involved in dopaminergic and GABAergic transmission, *Pcdh10*, linked to synaptic plasticity and social behavior (32), as well as *Nrxn3*, *Arhgap24*, respectively associated with addictive behavior (33, 34), major depressive disorder (35). In the hippocampus, *Ank3*, *Mgat5* and *Gbp6* were respectively found to be associated with bipolar disorder (36, 37), depressive-like state (38, 39), and autism spectrum disorder (40).

In addition, several sleep- and circadian-related genes were seen in features sets of behavioral variables. *Bhlhe40* (*Dec1*), and *Rorb* were identified as predictors of despair behavior in the hippocampus. *Crem*, which has been associated with the circadian regulation of melatonin synthesis and glucocorticoid secretion (41), was also a predictor of despair in the hypothalamus. Moreover, *Id2* and *Gtpbp1* contributing to the regulation of the circadian clock, as well as *Kcnj13* were hippocampal and hypothalamic predictors for anhedonia-like behavior.

Functional analysis of predictors underlying the corticosterone regulation identified several enriched pathways in the hippocampus. There were primarily associated with apoptosis and cell survival (e.g., role of Inhibitor of Apoptosis Proteins [IAP] in apoptosis, regulation of apoptosis by mitochondrial proteins, Apoptotic TNF-family pathways), stem cells, immune response (e.g., Interleukin 4 and 5 signaling pathways, Inducible COStimulator signaling pathway in T-helper cell), development (e.g., regulation of endothelial progenitor cell differentiation from adult stem cells), as well as several generic metabolic and signaling pathways (*SI Appendix*, Fig. S7).

### Pathways shared between REM sleep, corticosterone regulation and anhedonia

More than one third of the pathways associated with corticosterone regulation (37.5%) overlapped with pathways for REMS bout length and/or for EEG theta activity in the prefrontal cortex and hippocampus respectively (*SI Appendix*, Fig. S7). Most of these common pathways were associated with cell proliferation, differentiation and apoptosis. They included several apoptotic pathways involved in the extrinsic death receptor pathway (e.g., TNF receptor 1 signaling, FAS signaling cascade; apoptotic TNF-family pathways) and the intrinsic mitochondrial pathway (role of IAPs in apoptosis; regulation of apoptosis by mitochondrial proteins) (Table 1 and *SI Appendix*, Fig. S7).

In addition, REMS variables, and in particular REMS bout length and EEG theta activity also shared pathways identified with molecular correlates of anhedonia. These included pathways involved in cell proliferation and apoptosis (i.e., EGF-induced proliferation of Type C cells in subventricular zone [SVZ] of adult brain; cytoplasm/mitochondrial transport of proapoptotic proteins) and response to oxidative stress (NRF2 regulation of oxidative stress response). Molecular correlates of anhedonia were enriched in several pathways involved in development and stem cell processes (e.g., NOTCH1-mediated pathway for NF-κB activity modulation; EGFR signaling via small GTPases, FGFR signaling pathway, astrocyte and oligodendrocyte differentiation, regulation of transcription; *SI Appendix*, Fig. S7). Several enriched pathways were found for despair behavior, primarily in the hippocampus and included circadian rhythm process (*SI Appendix*, Fig. S7). None were common with REMS. Lastly, no overlap was observed between pathways associated with NREMS variables, corticosterone regulation and any of the investigated behavioral variables (*SI Appendix*, Fig. S8).

## Discussion

### REM sleep enhancement, a core adaptive response to chronic mild stress

The UCMS paradigm induced changes in physical, behavioral and neuroendocrine variables in accordance with previous reports (8, 21, 29). Here we also document the temporal emergence of physical, behavioral and neuroendocrine alterations and provide evidence that increase in REMS variables (i.e., 24-h duration, continuity, EEG theta oscillations) exhibited not only one of the largest effect sizes but were also the earliest response induced by stress. The longitudinal assessment of sleep throughout the UCMS paradigm furthermore demonstrated the differential effects of chronic stress on REMS and NREMS. The increased continuity and duration of REMS, and the increase in EEG theta activity during REMS, primarily reflecting hippocampal theta activity, imply that REMS is affected by UCMS in a positive manner. By contrast, NREMS continuity was decreased and EEG delta power in NREMS was not affected. While obtained in mice, the changes observed in sleep and their effect sizes agree well with meta-analyses performed in clinical depressive populations demonstrating that reduced sleep continuity and NREMS percentage showed smaller effect sizes than REMS (15, 18).

### Close association between REMS enhancement and the corticosterone regulation to UCMS

While REMS enhancement and alterations in the HPA axis negative feedback regulation of corticosterone have been previously reported in preclinical studies of chronic stress or stress-vulnerable rodents (9, 10, 29, 42), the current data demonstrate for the first time the very close association between these two phenotypes. Although the mechanisms underlying this association are unknown, one potential candidate is melanin-concentrating hormone (MCH). Substantive data suggest a role of MCH in the regulation of REMS. MCH-producing neurons are active during REMS (43) and intracerebroventricular or microinjection in the ventrolateral periaqueductal gray with MCH increased REMS in rats (44). Recent optogenetic studies confirmed the role of MCH in the promotion and maintenance of REMS (45). In addition, intracerebroventricular and microinjection of MCH stimulate the release of both adrenocorticotropic hormone and corticosterone (46).

In humans, increased REMS (47) and HPA axis dysregulation (14) were previously shown to correlate with remission and recovery in major depressive disorder. In this context, REMS has been proposed to play a central role in emotional processing and memory consolidation (48-51), with recent studies demonstrating a causal role for EEG theta activity during REMS in contextual and extinction memory consolidation, both in rodents and humans (52, 53). Furthermore, REMS is suppressed by most antidepressants (54) and under remarkably accurate homeostatic control which is modulated by some antidepressants (55).

### Transcripts and associated processes affected by 9-week exposure to chronic mild stress

Transcriptome changes, assessed by differential expression, were relatively small, which is consistent with previous reports despite the different methodologies employed in the various studies (21, 22). Also in line with previous reports, most changes were observed in the hippocampus (21, 22, 56), confirming its sensitivity to stress (57). Our data highlight that despite the tissue-specific differential expression of genes, in particular in the hypothalamus compared to the hippocampus and prefrontal cortex, many enriched biological processes were shared across brain regions. These include processes associated with inflammatory and immune responses, including innate immune responses, which had been reported in the hippocampal transcriptome following 13-day chronic social stress or several weeks of UCMS (19-21). Thus, our data highlight that para-inflammation appears to be a common mechanism in the three brain regions investigated, and is part of the protective adaptive response to chronic stress which is in line with a recent framework emphasizing that inflammatory signals contribute to restore homeostasis (58). This observation agrees well with the emerging view that chronic stress and stress-related diseases, such as major depression, share inflammation as a common mediator (59, 60).

Here we also identified that a number of DEGs following chronic stress were involved in sleep and circadian rhythms, with some cortical DEGs associated with sleep quality (*Cdhr1* (61)), sleep duration (*Ttr* (62)), entrainment of anticipatory behaviour (*Mc3r* (63)), or circadian rhythms (*Ppp1r3b* (64)). In the hippocampus, *Rorc* has been associated with circadian rhythms (65) and in the adaptive immune system (66). Nonetheless, the transcriptome changes identified after 9-week of chronic mild stress in the four tissues differed from the response to acute sleep deprivation in the brain transcriptome and identified only two transcripts (*Hspa1a, Hspa1b*) in blood that were previously shown to be induced by sleep loss independent of corticosterone and time of day (67). These heat shock proteins play a key role in a variety of cellular processes including protection of the proteome during cellular stress. Furthermore, the blood transcriptome showed a unique signature with many biological processes involved in cellular stress, such as oxidative stress and unfolded protein response. These data highlight the value of using the blood transcriptome in animal studies to identify biomarkers in response to challenges and draw comparison with patient populations (22). Several DEGs encoding neuropeptides (*Pmch, Vip, Avp*) may provide a link at the neuroendocrine level between the stress-induced REMS alterations and the HPA axis dysregulation. *Pmch* encodes the precursor of MCH and neuropeptide-glutamic acid-isoleucine (NEI). These co-expressed neuropeptides have been involved in stress-induced release of adrenocorticotropic hormone (68). As discussed above, several lines of evidence suggest a role of MCH in REMS and a recent study showed that intracerebroventricular or microinjection of NEI in the ventrolateral periaqueductal gray also increased REMS in rats (44). In addition, vasoactive intestinal peptide (VIP) and vasopressin have also been implicated in REMS regulation and sleep continuity (69-71). While VIP has been involved in the circadian release of glucocorticoids (72), vasopressin play a role in HPA axis activity (73), stress coping and social behavior (74).

### REM and NREM sleep, circadian rhythmicity, corticosterone regulation, anhedonia and transcriptomic predictors identified using machine learning

Transcriptomic predictor sets were overall relatively small in accordance with a previous study in which phenotypes were grouped into categories (75). Application of a machine learning approach which controlled for the unspecific effects of UCMS demonstrated that NREMS and REMS shared very few predictors, further emphasizing the contrast between these two sleep states observed at the electrophysiological level in this study. Furthermore, no overlap was observed between pathways associated with predictors for NREMS variables, corticosterone regulation or behavioral phenotypes.

Machine learning identified a number of circadian-related transcripts as predictors of NREMS variables, as well as despair and anhedonia-like behaviors. This is consistent with a recent study correlating UCMS-induced depressive-like behavior with circadian rhythm alterations in brain tissues (76), and the growing recognition that circadian rhythmicity may play a role in mood regulation and development of mood disorders (77, 78). It should be noted that although we observed changes in sleep, at the behavioural level circadian rhythmicity was not much affected. Circadian changes were limited to an increase in wakefulness and activity during the light phase and a reduction of wakefulness and activity during the dark phase, which may be interpreted as a reduction in circadian amplitude.

Hippocampal transcriptomic predictors of 24-h REMS duration were associated with pathways involving stem cells differentiation and hedgehog signaling. Inhibition of hedgehog signaling by glucocorticoid treatment has been shown to decrease hippocampal cell differentiation (79). In addition, several enriched pathways in apoptosis and cell survival were among the molecular signatures characterizing REMS continuity variables and EEG theta power, in the cortex and hippocampus respectively. The identified predictors and related pathways specifically associated with REM in the cortex, a brain region not necessarily implicated in the generation of REMS. These predictors may therefore reflect effector system by which REMS exerts its adaptive response to chronic stress. In this context, the endoplasmic reticulum (ER) stress response pathway was common to REMS continuity and theta power, suggesting that the ER stress response is part of the adaptive response to chronic stress. ER stress, which may lead to apoptosis (80), has been recently shown to be induced during social isolation in *Drosophila* (81). The link between REMS, cell proliferation and apoptosis has been primarily studied in the context of neurogenesis following REMS loss (82-85).

We identified overlapping pathways between the corticosterone regulation and REMS continuity variables and/or theta power, which were primarily involved in apoptosis and cell survival. These included several members of the Tumor-Necrosis-Factor signaling (e.g., TNFR1; Apo-L(TNFSF10)) which triggers a broad spectrum of actions at the cellular level, as well as a number of processes playing a role in the mitochondrial intrinsic pathway, such as the inhibitors of apoptosis family of proteins.

Here we also identify for the first time shared molecular pathways underlying the interindividual variation in anhedonia, a core symptom of depression, with REMS continuity and theta power in response to chronic stress. They included pathways playing a role in oxidative stress or apoptosis with the pathway involved in the transport of proapoptic proteins linking the Jun amino-terminal kinases (JNK) signal transduction pathway and the mitochondrial apoptotic machinery (86). A causal role for mitochondrial genes was recently proposed as part of the processes in the striatum linking REMS parameters and stress-induced anxiety-like phenotype (75) and the contribution of mitochondrial dysfunction in major depression is emerging (87).

### Limitations

Limitations of this study include a relatively small sample size and a conservative choice of setting the statistical significance at FDR-adjusted *P* < 0.05, which may have led to an underreporting of significant effects. Another limitation relates to experimental constraints which precluded an assessment of the temporal association between behavioral phenotypes and transcriptomic changes. Nevertheless, the high intra-individual stability of many of the phenotypes indicates that the observed transcriptomic changes at the end of the experiment are relevant to the phenotypes throughout the UCMS. A final limitation is that this study was only conducted in male mice. Hypotheses based on the current data may be tested in future studies in which sex differences in sleep disturbances and their underlying molecular mechanisms in the context of chronic stress could be investigated (88).

### Conclusion

This study provides a comprehensive characterization of sleep changes induced by chronic stress, with REMS increase being the earliest marker of a stress response. Alteration in corticosterone regulation and REMS have both been implicated in the response to emotional experiences. Our data show that apoptosis processes including mitochondrial pathways are associated interindividual variation in REMS, changes in corticosterone regulation and anhedonia and may mediate adaptation to waking experiences.

## Materials and Methods

### Animals

Male BALB/cJ mice (*n* = 18; B&K Universal Ltd, Grimston, Aldbrough, Hull, UK) were housed in groups prior to the start of the experiment, under a 12-h light/dark cycle, controlled ambient temperature (20-22°C) and relative humidity (55 ± 10%) with food and water *ad libitum*. Experimental procedures were approved by the University of Surrey Animal Welfare and Ethical Review Body and were carried out in accordance with the UK Animals (Scientific Procedures) Act 1986.

### Surgery and experimental setup

Mice aged 12.4 ± 1.4 (Mean ± SD) weeks were implanted with a telemetry transmitter (TL11M2-F20-EET, Data Sciences International, St. Paul, MN, USA) connected to electrodes for continuous EEG/EMG recordings (see *SI Appendix* for details), as previously described (89). After surgery, animals were allowed to recover for 5.6 ± 1.4 weeks (Mean ± SD). Mice were singly housed and randomly assigned to the control and the UCMS groups (*n* = 9 per group). Home cages were placed in ventilated cabinets with a light intensity of 200 ± 13.4 mW/m2 (~60 lux) at the level of the cage bottom during the light period with a 12-h light-dark cycle. Baseline data collection started when the mice were 18 weeks of age with baseline measurements, before the start of the UCMS protocol.

### Unpredictable chronic mild stress (UCMS) paradigm

Briefly, mice were daily subjected to various socio-environmental low intensity stressors (social stress, cage tilting, confinement stress, cage without sawdust, soiled sawdust, damp sawdust, sawdust change, water stress and predator [rat] odor) according to an unpredictable schedule for nine weeks (28) (see *SI Appendix* for details; Fig. 1A and *SI Appendix*, Table S1). No stressor was applied during the light period.

### Physical and corticosterone regulation assessments

Body weight and coat state were assessed weekly and twice a week, respectively, as markers of the progression of the UCMS-evoked syndrome (28). The coat state, an index of self-care behavior, was assessed as previously described (28) (see *SI Appendix* for details). Baseline values of the coat state and the body weight were assessed 3 days before the beginning of the UCMS paradigm. To assess the effects of UCMS on the HPA axis negative feedback-regulated corticosterone, the dexamethasone (DEX) suppression test was used as previously described with minor modifications (29) (see *SI Appendix* for details).

### Sleep recordings and analyses

EEG/EMG signals were recorded continuously for 48-h at several time points throughout the UCMS protocol (see *SI Appendix* for details). The data analyzed consisted of 24-h recordings starting at dark onset (ZT12). Vigilance states of 10-s epochs were determined using SCORE-2004™, an automated sleep scoring system based on an updated version of the real-time sleep/wake monitoring system SCORE™ (90). EEG power spectra were computed for consecutive 10-s epochs by a fast Fourier transform and determined for NREMS, REMS, and wakefulness and expressed as a percentage of total EEG power (see *SI Appendix* for details). EEG delta power during NREMS and EEG theta power during REMS were computed by adding the EEG power in the frequencies ranging from 1 to 4.5 Hz and from 6 to 9 Hz, respectively. Mean values were individually normalized to the mean total power in NREMS and REMS.

### Circadian locomotor activity

Locomotor activity was measured with a passive infrared motion detection sensor positioned on the cage lid. Activity counts were recorded continuously using ClockLab (Actimetrics, Wilmette, IL, USA). Averaged daily activity for the 12-h light and dark periods were computed and analyzed per week using Matlab 2016a (MathWorks, Natick, MA, USA).

### Behavioral testing

Behavioral tasks were performed as previously described to assess self-centered behavior (grooming test), motivation (nest building test), anhedonia (reward-driven exploratory test), social preference (social novelty preference test), aggressiveness (resident-intruder test), anxiety (novelty-suppressed feeding test) and despair behavior (forced swim test) (28, 29, 91, 92) (see *SI Appendix* for details). All behavioral tests were performed between Zeitgeber time (ZT) 13 and ZT17 (i.e., during the dark, active phase) under red light, with a maximum of one test per day.

### RNA extraction, library generation and sequencing

At the end of the protocol, mice were euthanized between ZT12.5 and ZT14.5. Total RNA was extracted from dissected prefrontal cortex, hippocampus, hypothalamus and whole blood samples. RNA quantity and integrity were assessed and total RNA libraries were prepared using 100 ng of total RNA from each sample. One-hundred base pair paired-end reads were generated for each library using the Illumina HiSeq®2500 (Illumina Inc., San Diego, CA, USA). At least 40 million reads were obtained for brain tissue samples, whereas for the blood samples a minimum of 60 million reads was obtained (for additional details, see *SI Appendix*).

### Transcriptome analysis

Quality control (QC) of sequencing reads, read alignment and feature quantification were performed using a RNA-Sequencing workflow developed by Eli Lilly and Company (for additional details, see *SI Appendix*).

#### Rank Products analysis for DEG identification

Differential expression analysis between control and UCMS-subjected animals was performed using the non-parametric Rank Product statistical method that is independent of inter-class variability (93), using the R Bioconductor package RankProd (94). Significance was set at a proportion of false positive (PFP; abbreviated *P*_pfp_ in the text) < 0.05.

#### Selecting genes associated with a given phenotype

To robustly select relevant predictors, we used a form of penalized regression referred to as Elastic net. Elastic-net learning was performed using the R package glmnet (95). Analyses were focused on sleep variables, three behavioral variables and corticosterone regulation, the values of which were z-scored within group in order to identify associations with phenotypes independent of any ‘unspecific’ group effect. We only report predictor sets for variables achieving a positive Pearson correlation r between observed and cross-validated prediction values > 0.31.

#### Functional annotation and interpretation

Lists of genes associated with an attribute, either by Rank Products analysis or Elastic-net learning, were subjected to Gene Ontology (GO) enrichment analyses (Gene Ontology (GO) processes and/or Pathway maps) using MetaCore™ (GeneGo, Thomson Reuters; https://portal.genego.com/; updated mid-June 2018). Significant enrichment was defined by nominal *P*-value (*P*_nom_) < 0.05 and False Discovery Rate-adjusted *P* (*P*_adj_) < 0.05).

#### Similarity of gene lists between variables within a tissue

The hypergeometric test was used (phyper function in R) to generate a ‘raw’ nominal *P*-value (*P*_nom_). The resultant P-value was based on the observed overlap between the two gene lists, the size of each gene list, and the size of the tissue-specific transcriptome as background. All raw *P*-values within a tissue were adjusted (abbreviated to *P*adj in the text) using the Benjamini and Hochberg false discovery rate correction (p.adjust function in R).

### Statistics on sleep, other behavioral, physical and endocrine variables

Unless otherwise stated, data were analyzed with SAS 9.2 (SAS Institute, Cary, NC, USA). For repeated measures, data were analyzed as dependent variables in a general linear mixed model using PROC MIXED for analysis of variance (ANOVA) with group (treatment: UCMS *versus* control) and time (day, treated as repeated measure with spatial power anisotropic variance-covariance matrix) as categorical explanatory variables with baseline as a covariate (no group effect was found at baseline for all measures). *Post-hoc* multiple pair-wise comparisons (contrast between UCMS and control groups) were assessed using the ESTIMATE option of PROC MIXED. Output data are expressed as least squares means (LSmeans) with 95% Confidence Intervals (CIs). For non-repeated measures, PROC TTEST for pair-wise comparisons was used using Pooled or Satterthwaite methods for equal and unequal variances, respectively. Output data are expressed as mean ± standard error of the mean (SEM). To visualize statistical effect sizes, the Cohen’s *f*^2^ effect sizes were calculated (96). To assess the variability within experimental groups throughout time, intra-class correlations (ICC) were also computed (30). Kendall’s partial correlations (Kendall’s tau coefficients), with experimental group as the controlled variable, were computed using PROC CORR for generating the phenotypic associations between all output measures, while the false discovery rates (FDRs) using the Benjamini-Hochberg procedure for multiple testing correction were computed using the p.adjust function in R. Correlations were considered significant at FDR-adjusted *P*-value (*P*_adj_) < 0.05 (nominal *P*-value: *P*_nom_). For repeated measures, the average of the last three measures was used for the calculation of the correlation table. Scoring agreement among raters (97) was assessed using Kendall’s coefficient of concordance *W* computed in R using the irr package (https://cran.r-project.org/web/packages/irr/index.html).

## Acknowledgements

We thank Drs. Sig Johnsen and Jeewaka Mendis for advice and support with statistical analyses, Drs. Daan van der Veen, Bruno Martynhak for help with locomotor activity data analyses, Dr. Gillian Stenson for support during data acquisition and double-scoring of behavioral tasks. We thank Drs. Hugh Marston and Dale Edgar for support, Dr. Lisiane Meira for helpful discussions, and Dr. Simon Archer for comments on the manuscript.

**Author contributions:** M.N., K.A.W., D.-J.D. and R.W.-S. designed research; M.N. performed research; M.N., A.P.McC., H.W., C.S.M.-L., E.E.L., K.M. and N.L. contributed new reagents or analytic tools; M.N., H.H., A.P.McC., H.W., C.S.M.-L., E.E.L., K.M., N.L., D.-J.D. and R.W.-S. analyzed data; and M.N., K.A.W., D.-J.D. and R.W.-S. wrote the paper.

